# Transcriptional and post-transcriptional regulation of young genes in plants

**DOI:** 10.1101/2021.12.20.473517

**Authors:** Vivek Kumar Raxwal, Somya Singh, Manu Agarwal, Karel Riha

**Author notes:** These authors contributed equally (Vivek Kumar Raxwal, Somya Singh).

## Abstract

New genes continuously emerge from non-coding DNA or by diverging from existing genes, but most of them are rapidly lost and only a few become fixed within the population. We hypothesized that young genes are subject to transcriptional and post-transcriptional regulation to limit their expression and minimize their exposure to purifying selection. We found that young genes in rice have relatively low expression levels, which can be attributed to distal enhancers, and closed chromatin conformation at their transcription start sites (TSS). The chromatin in TSS regions can be re-modeled in response to abiotic stress, indicating conditional expression of young genes. Furthermore, transcripts of young genes in Arabidopsis tend to be targeted by nonsense-mediated RNA decay, presenting another layer of regulation limiting their expression. Together, these data suggest that transcriptional and post-transcriptional mechanisms contribute to the conditional expression of young genes, which may alleviate purging selection while providing an opportunity for phenotypic exposure and functionalization.

## Introduction

The advent of whole-genome sequencing led to the discovery of a subset of new genes in all domains of life that lack homologs in other lineages (Dujon 1996; Fischer and Eisenberg 1999; Arendsee, et al. 2014; Stein, et al. 2018). These genes, also called orphan or evolutionary young genes, may arise from pre-existing genes by diverging until no homology remains, or they may be born *de-novo* from non-coding DNA (Tautz and Domazet-Loso 2011; Rodelsperger, et al. 2019; Van Oss and Carvunis 2019). In contrast to old genes, young genes are short, rapidly evolving, and usually do not have essential functions. They are therefore under weaker positive selection (Wolf, et al. 2009; Cai and Petrov 2010). Furthermore, production of nonfunctional proteins from young genes may represent an energetic burden for the cell, and their evolutionarily non-optimized structure can lead to nonproductive interactions, some of which may interfere with cellular functions (Drummond, et al. 2005; Drummond and Wilke 2008). Consequently, despite increasing the population’s genetic diversity, most young genes are rapidly lost either due to genetic drift or purging selection, and only a few are fixed in the genome (Palmieri, et al. 2014). This raises the question of whether some young genes have the means to hide from purifying selection, expanding their lifespan in a genome and thereby increasing their chances of acquiring novel functions.

One possibility to mitigate the effect of purifying selection is by limiting the extent of expression and/or conditioning it by developmental or environmental cues. Indeed, young genes often have low expression due to the deposition of repressive chromatin marks and the lack of well-developed cis-regulatory elements required for transcription (Donoghue, et al. 2011; Werner, et al. 2018). The low expression level of young genes can lessen the burden of protein misfolding, hence reducing the negative selection pressure (Drummond, et al. 2005; Drummond and Wilke 2008). Further, spatial or temporal restriction of expression may expand the lifespan of young genes by avoiding untimely exposure to natural selection and provide an opportunity for phenotypic manifestation and functionalization. In nematodes, young genes are born in the vicinity of enhancers and utilize their cis-regulatory elements to express themselves in limited cell-types and tissues (Majic and Payne 2020). Moreover, the permissive transcription environment of isolated compartments, such as the testes in animals and pollen grains in plants, provides a perfect breeding ground for young genes to mature and gain function (Vinckenbosch, et al. 2006; Cui, et al. 2015).

In this study, we investigated the mechanisms that contribute to the low or conditional expression of young genes via transcriptional and post-transcriptional regulation in rice and Arabidopsis. We propose that the restricted expression of young genes through these mechanisms can mitigate exposure to negative selection and offer an opportunity for phenotypic manifestation under certain conditions, which permits gene functionalization and genetic fixation in a population.

## Result and Discussion

### Young genes are associated with closed chromatin and distal enhancers

We performed a protein-based homology search across the tree of life to determine the evolutionary age of protein-coding genes present in the rice genome using Phylostratr (Arendsee, et al. 2019). The genes were grouped into phylostrata (PS) according to their evolutionary age such that PS1 contains genes with the oldest known ancestor homolog, whereas the last phylostrata (PS19) includes the evolutionary youngest genes with no known ancestral homolog (**Supplementary Figure 1**). To understand differences in gene regulation and the underlying reasons, we compared the steady-state expression of evolutionary old (PS1) and young genes (PS19) and observed that young genes largely had low expression relative to the old genes (**Figure 1A**).

**Figure 1:**
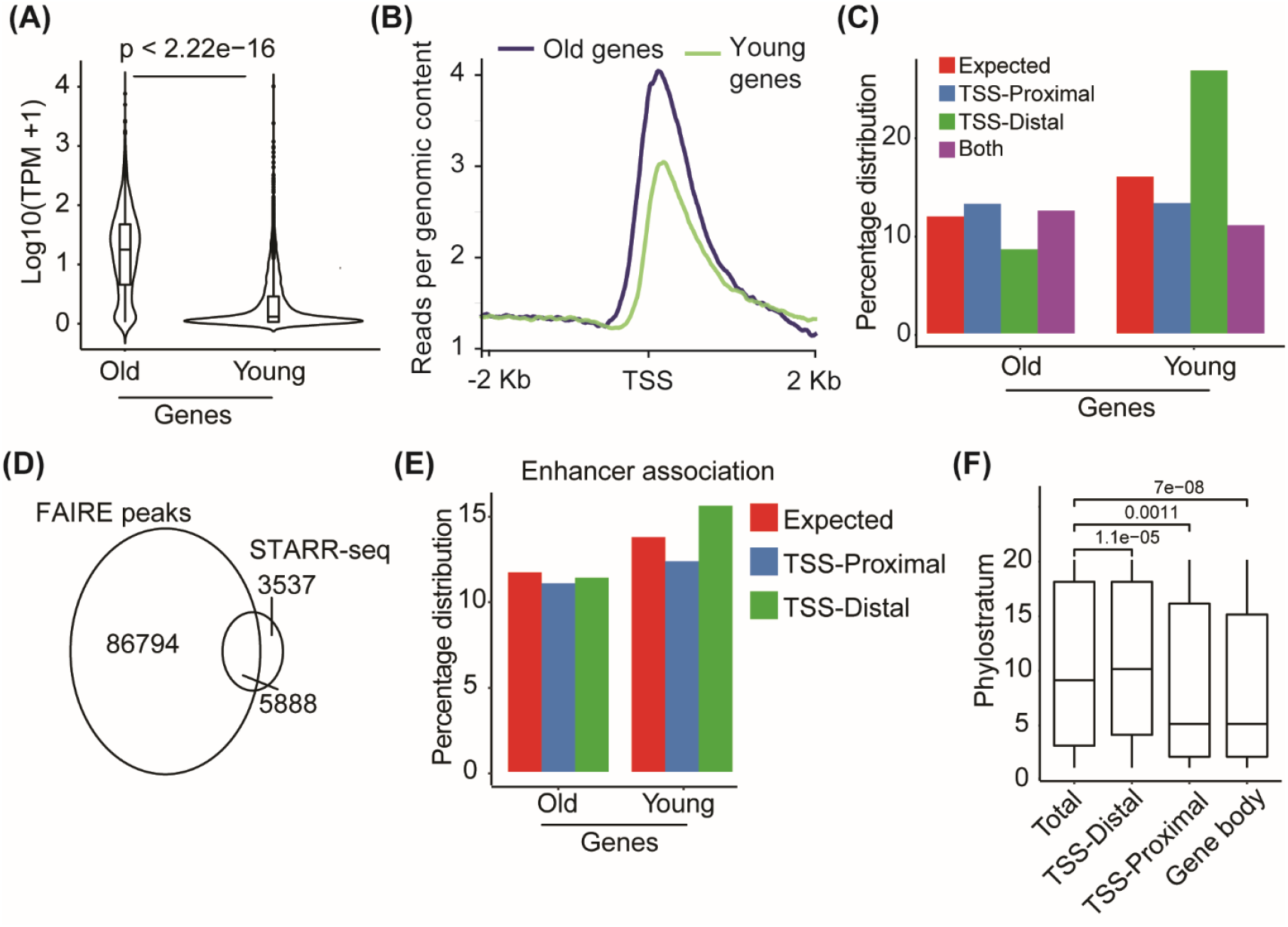
Transcriptional regulation of the young genes. (A) A violin plot showing the expression of old and young genes. The expression of genes in TPM (transcript per million) was log10 transformed before plotting. The statistical significance of the difference was calculated using the Mann-Whitney test. (B) A line plot showing normalized coverage of FAIRE-seq reads around transcription start site (TSS) of old (PS1) and young (PS19) genes. (C) A bar plot was drawn to show the enrichment of FAIRE-seq peaks at either the TSS-proximal (up to 1.5 kb upstream of TSS), TSS-distal (> -1.5 kb from TSS), or both (FAIRE-seq peak present at TSS-proximal as well as TSS-distal intergenic region) of old and young genes. Enrichment was higher than background (expected), and was calculated as the total percentage of genes present in the old or young gene category. (D) An area proportional Venn diagram showing overlap of FAIRE-seq identified open chromatin regions and STARR-seq identified enhancers. (E) A bar plot was drawn to show enrichment of enhancers at the TSS-proximal (up to 1.5kb upstream of TSS) and TSS-distal (> -1.5kb from TSS) regions of old and young genes. The enrichment was seen over the background (expected), calculated as the total percentage of enhancers present in the old or young gene category. (F) A box plot showing the association of STARR-seq identified enhancers with the age of genes. Enhancer is characterized as TSS-Distal if it is present > 1.5kb upstream of the TSS, whereas TSS-Proximal enhancers are located within 1.5kb upstream of TSS. If an enhancer is present at any part of the gene body, then it is characterized as Gene body enhancer. The age of the nearest gene is plotted at the Y-axis, where phylostratum 1 denotes the oldest and phylostratum 19 denote the youngest class of genes.

Young genes have been reported to possess heterochromatic epigenetic signatures and a closed chromatin state in Drosophila, nematodes, and Arabidopsis (Li, et al. 2016; Werner, et al. 2018; Zhang and Zhou 2019). To investigate whether the lower expression of young genes in rice was due to the closed chromatin architecture of their regulatory regions, we employed Formaldehyde Assisted Isolation of Regulatory Elements (FAIRE-seq). We observed that the TSS-proximal regions of the young genes have a more closed chromatin conformation relative to old genes (**Figure 1B**). This is in agreement with higher levels of repressive histone modifications (H3K9me2 and H3K9me1) and lower levels of permissive modifications (H3K9ac and H3K4ac) at the TSS of young genes compared to old genes (**Supplementary Figure 2**). These results suggest that a generally closed chromatin conformation limits the expression of young genes in rice.

Interestingly, our FAIRE-seq analysis in rice revealed an enrichment of open chromatin at the distal intergenic regions (> 1500 bp upstream from TSS) of young genes (**Figure 1C**), which has also been observed in Arabidopsis (**Supplementary Figure 3**). Open chromatin at a distal intergenic region is indicative of an enhancer (Zhu, et al. 2015), and enhancers have been suggested to be involved in the birth of young genes (Werner, et al. 2018; Majic and Payne 2020). To determine whether distal upstream regions of young genes indeed contain enhancers, we determined the overlap between open chromatin, which we identified by FAIRE-seq, and enhancers, identified by STARR-seq, in rice (Sun, et al. 2019). Our overlap analysis revealed that more than 60% of the known enhancers overlapped with open chromatin regions (**Figure 1D**). Next, we compared the enrichment of enhancers in TSS-proximal and TSS-distal regions. We found that enhancers are enriched in the TSS-distal regions of young genes but not in the TSS-distal regions of old genes (**Figure 1E**). We also observed that the average age of the nearest gene to distal intergenic enhancers is younger in contrast to the enhancers present in the TSS-proximal region (**Figure 1F**). Interestingly, we did not detect an enrichment of enhancers in TSS-proximal regions of young genes. This is in contrast to previous findings in nematodes which suggested that genes are born within open regions of enhancers (Werner, et al. 2018). Instead, our result suggests that young genes tend to associate with distally positioned enhancers, possibly because existing cis-regulatory elements are able to restrict their expression (Spitz and Furlong 2012; Andersson, et al. 2014). The utilization of distal enhancers likely contributes to the lower expression observed for young genes, which reduces the cost to the cell of translating misfolded and possibly toxic proteins. Moreover, enhancers evolve faster than promoters, thereby providing an opportunity for young genes to evolve together with enhancers as they acquire novel transcription regulatory networks over time (Prud’homme, et al. 2007; Villar, et al. 2015).

### Abiotic stress alters chromatin architecture at TSS of young genes

Expression of young genes can provide species-specific adaptation to environmental challenges (Donoghue, et al. 2011; Carvunis, et al. 2012), suggesting that their chromatin architecture and transcription are responsive to external stimuli. Since many young genes in rice have a closed chromatin conformation in the promoter-proximal regions, we investigated whether abiotic stresses have the potential to affect chromatin conformation. We performed FAIRE-seq on rice seedlings subjected to varying durations of cold, heat, and salt stress (**see methods**). Except for cold stress (12 h), all of the abiotic stresses we tested increased chromatin accessibility around the TSS of young genes (**Figure 2**). These results suggest that chromatin architecture can be re-modeled upon exposure to external environmental factors, allowing young genes to gradually evolve interactions between cis-regulatory elements and regulatory proteins, thereby providing an opportunity to increase their expression and gain function.

**Figure 2:**
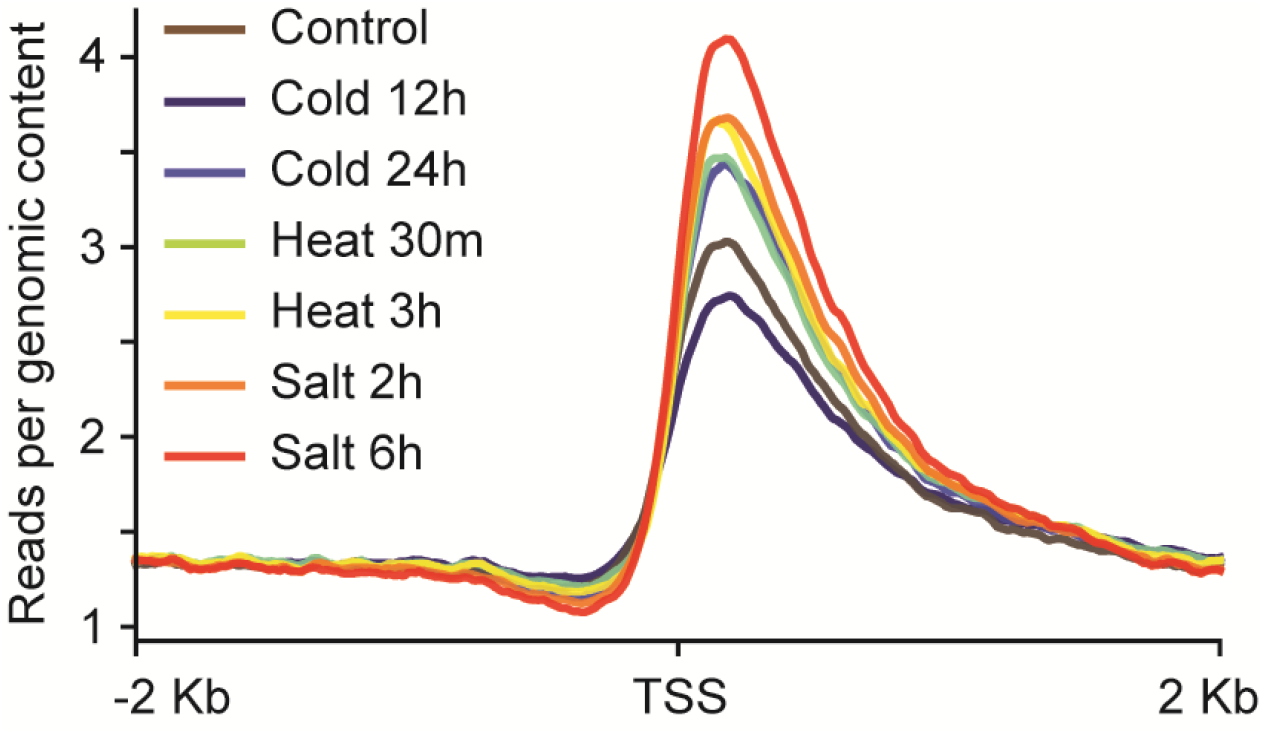
Abiotic stress re-models the chromatin architecture of young genes. Normalized coverage of FAIRE-seq reads around the TSS of young genes plotted as a line plot.

### Young gene transcripts are targeted by nonsense-mediated RNA decay

Apart from transcriptional regulation, the expression of young genes can be reduced by post-transcriptional regulation. Young genes in Drosophila have been associated with increased occurrence of premature translation termination codons (PTC) (Yang, et al. 2015). Since PTC-containing transcripts are degraded by the non-sense mediated RNA decay (NMD) pathway (Lykke-Andersen and Jensen 2015), we hypothesized that expression of young genes is affected by NMD. To examine this hypothesis, we took advantage of genetic and genomic resources available in Arabidopsis (Raxwal, Simpson, et al. 2020). Indeed, compared to older genes, young genes were enriched for PTCs and long 3`UTRs, both hallmark NMD features (**Figure 3a)**. We also observed an increased incidence of PTCs in young genes of maize and Arabidopsis (**Supplementary Figure 4**). We further found that young genes have significantly reduced transcript stability compared to old genes (**Figure 3b**). The Arabidopsis *upf1 pad4* mutant, which is severely compromised in NMD (Riehs-Kearnan, et al. 2012; Raxwal, Simpson, et al. 2020), exhibits a more pronounced increase in the expression of young genes than old genes (**Figure 3c**). These results suggest that young genes are subject to post-transcriptional regulation by NMD.

**Figure 3:**
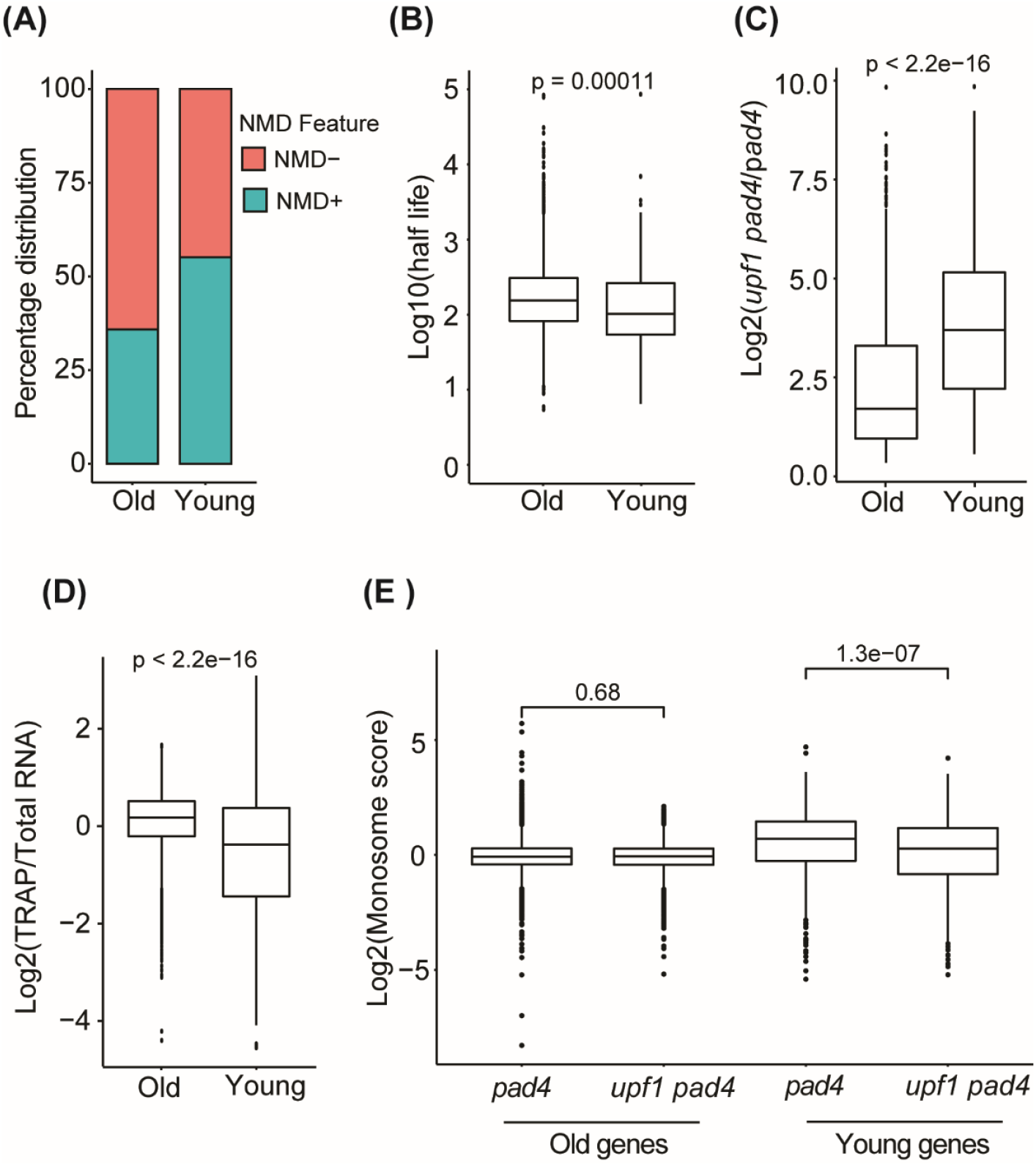
Nonsense-mediated RNA decay (NMD) post-transcriptionally regulates the abundance of young genes. (A) The accumulation of NMD features in old and young genes is presented as a bar plot. (B) A box plot representing half-life (log10 transformed) of the old and young genes. (C) The relative change in expression of old and young genes due to UPF1 deficiency. (D) The association of old and young genes with the ribosome is shown as a box-plot. (E) A box-plot depicting association of old and young genes with monosome and polysome either in *pad4* or UPF1-null (*upf1 pad4*).

The increased targeting of young gene mRNAs by NMD implies lower levels of translation. Therefore, we next evaluated the relative efficiency of translation by scoring differences in ribosome association between old and young genes using the Targeted Purification of Polysomal mRNA (TRAP) dataset (Chantarachot, et al. 2020). We observed that transcripts from older genes had substantially higher levels of ribosome association than transcripts from younger genes, indicating less efficient translation of young genes (**Figure 3d**). Previously, we determined that inactivation of NMD leads to increased translation of NMD-targeted transcripts, which was manifested as a shift of the transcripts from monosomal to the polysomal fraction of ribosomes (Raxwal, Simpson, et al. 2020). If NMD contributes to the translational repression of young genes, we also anticipated a similar trend for young gene transcripts. Indeed, NMD inactivation led to a significant decrease of the monosome score for young transcripts, which indicates their shift to polysomes and increased translation (**Figure 3E**). In contrast, no significant difference was observed in the monosome score of older genes in the presence or absence of UPF1 (**Figure 3E**). These results further substantiate the role of NMD in limiting expression of young genes at the level of RNA stability and translation. Because NMD efficiency changes in response to environmental and developmental cues (Hug, et al. 2016; Raxwal and Riha 2016), the repression of young genes by NMD is not constitutive. Rather, it could be lifted under certain conditions. The observation that young genes are subject to regulation by NMD is in line with our hypothesis where we proposed that NMD can act as an evolutionary capacitor, permitting accumulation of cryptic genetic variation and expose it conditionally to the selection (Raxwal and Riha 2016).

In conclusion, we propose that young genes tend to have low levels of expression due to closed chromatin, basal transcription by enhancers, and post-transcriptional degradation by NMD to avoid their untimely exposure to purifying selection. All three of these mechanisms are responsive to environmental stress and developmental signals, which provide an opportunity for the expression and phenotypic manifestation of young genes; a prerequisite for gene functionalization.

## Methods

### Plant material, growth conditions, and stress treatment

*Oryza sativa* L. Japonica nipponbare was grown in a plant growth chamber (Conviron®) at 28°C under a photoperiod of 16 h/8 h. 14-day old seedlings were subjected to heat (42°C for 30 min and 3 h), cold (4°C for 12 h and 24 h), and salt (250 mM NaCl for 2 h and 6 h) stress. The 14-day old seedlings grown at 28°C were taken as the control sample.

### Formaldehyde Assisted Isolation of Regulatory Elements (FAIRE)-seq

FAIRE-seq was performed on two independent replicates of 14-day old stress-treated and control seedlings as per the protocol described previously (Raxwal, Ghosh, et al. 2020). The raw sequencing reads were aligned to the Rice reference genome (IRGSP 1.0) using Bowtie2 with default parameters (Langmead and Salzberg 2012). The reads aligning to the region of the chromosome with a known insertion site of the mitochondrial and chloroplast genome were removed. To remove reads mapped to multiple positions on the genome, reads with a mapping score of less than 10 were filtered out. To remove potential PCR duplicates, reads with the same start and end position were considered only once. Broad peaks were called by MACS2 with default parameters except the no-model option to identify open chromatin regions in the Rice genome (Zhang, et al. 2008). Further, peaks having read count <1 RPM in any biological replicates were removed from the analysis. A Scatter plot was generated to observe the reproducibility of peaks among biological replicates. The overlap analysis was performed using the R package ChIPpeakanno (Zhu, et al. 2010).

### RNA-seq and analysis

RNA-seq libraries from RNA extracted (the Qiagen plant RNA extraction kit) from two independent replicates were prepared using the TruSeq RNA sample preparation kit (Illumina Inc., USA). The libraries were sequenced for 50 bp single-end sequencing on Illumina’s HiSeq 2000 platform. The sequencing reads were filtered for quality using Trimgalore (https://www.bioinformatics.babraham.ac.uk/projects/trim_galore/) and high-quality reads were pseudo aligned to Rice transcriptome (ensemble 46 version) using Kallisto version 0.46.0 with default parameters (Bray, et al. 2016). The differential expression analysis was performed using a limma-voom pipeline enabled in 3D-RNA-seq (Guo, et al. 2021).

### Evolutionary age classification of genes, NMD features and mRNA half-life analysis

The evolutionary age or phylostratum of each peptide present in the Rice (Oryza sativa Nipponbare version), Maize (B73 RefGen_v4) and Arabidopsis (Araport 11) was determined using Phylostratr (Arendsee, et al. 2019). A transcript is defined as having an NMD feature if it has premature termination codon (PTC) before (greater than 50 bp) the last exon junction complex (EJC) or the length of its 3`UTR exceeds 350 bp. The mRNA half-life, TRAP, and NMD data were taken from (Sorenson, et al. 2018; Chantarachot, et al. 2020; Raxwal, Simpson, et al. 2020).

## Acknowledgment

This work was supported by the Ministry of Education, Youth, and Sports of the Czech Republic, European Regional Development Fund-Project REMAP (grant CZ.02.1.01/0.0/0.0/15_003/0000479 to K.R.), Faculty Research Program Grant from IOE, University of Delhi (grant IoE/2021/12/FRP to MA), India and PURSE grant from the Department of Science and Technology (grant to MA), India. VKR and SS are grateful for research fellowships from the Council of Scientific and Industrial Research (CSIR), India and Department of Biotechnology, India, respectively. We thank Dr Sridhar Sivasubbu and Dr Shamsudheen from the Institute of Genomics and Integrative Biology (IGIB), Delhi, India for helping with the sequencing.

## Supplementary figures

**Supplementary Figure 1.**
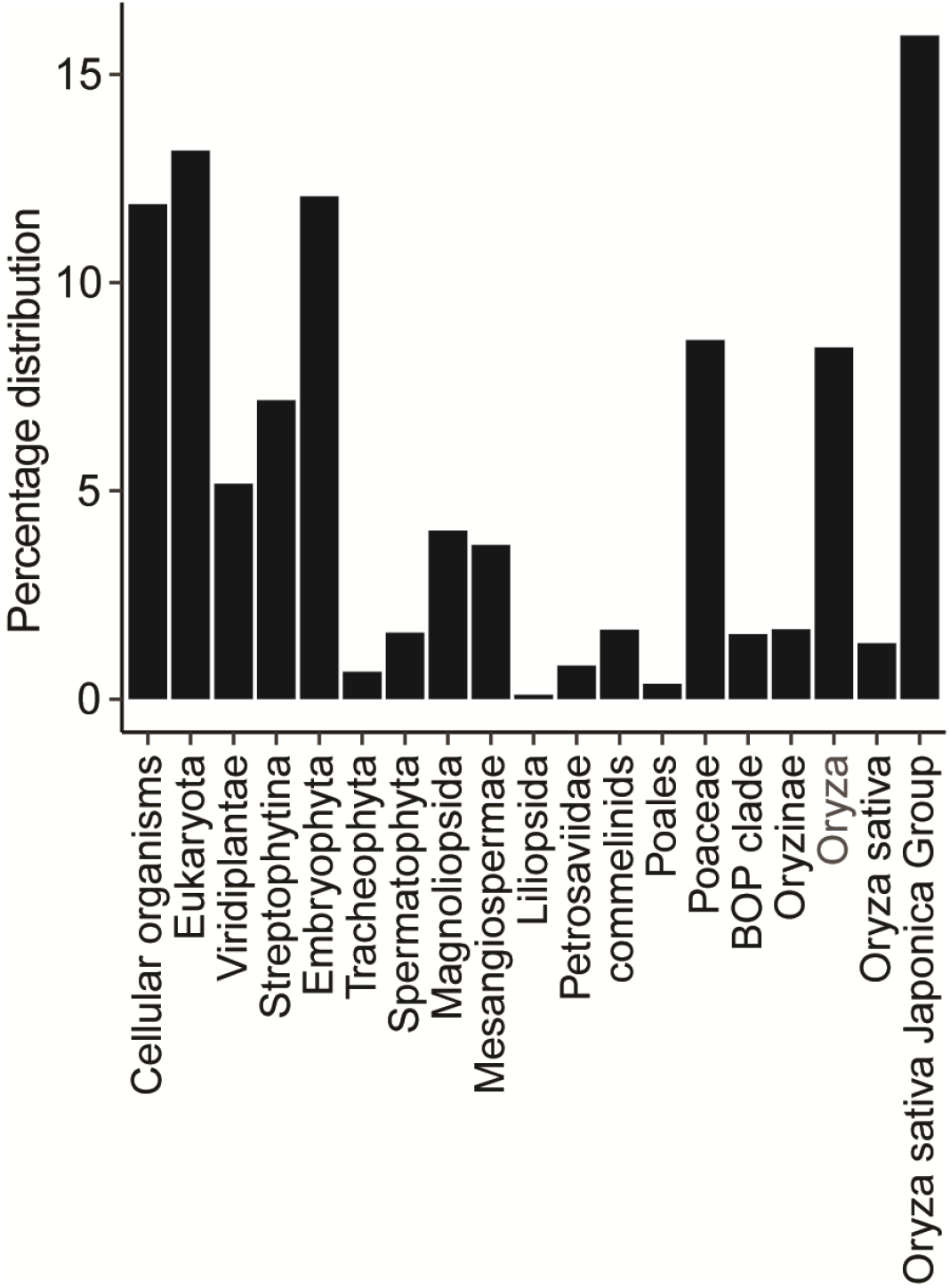
Phyllostratographic classification of rice genes showing the distribution of genes with respect to their evolutionary age.

**Supplementary Figure 2.**
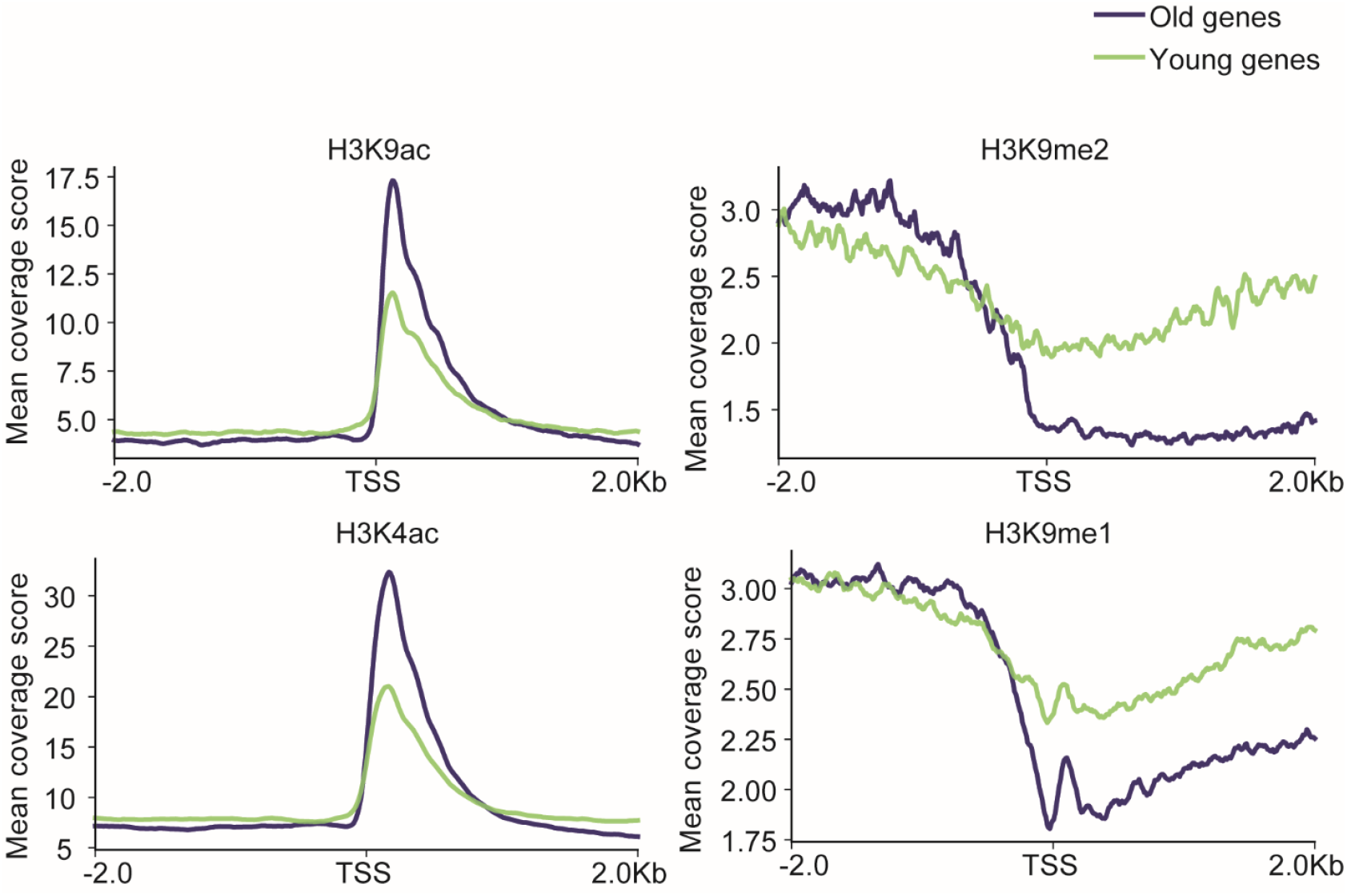
A line plot showing the deposition of various histone modification marks over the TSS of young and old genes.

**Supplementary Figure 3.**
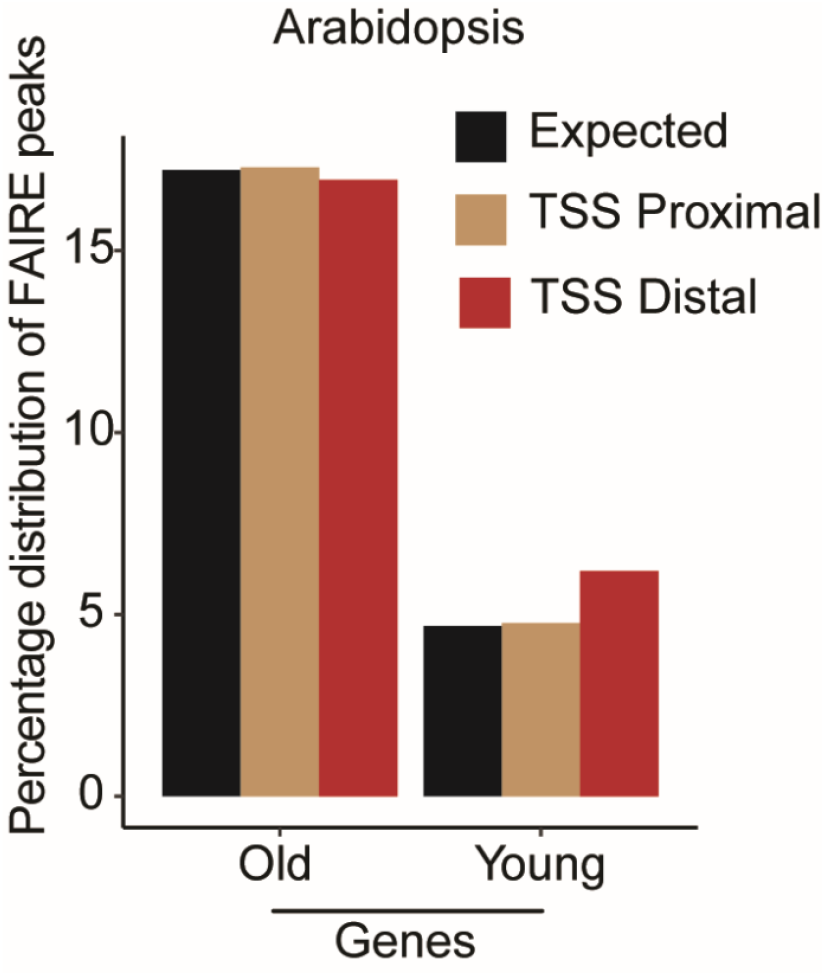
A bar plot showing the percentage distribution of FAIRE-peaks in either the TSS proximal (<-1.5kb of TSS) or TSS distal region (> -1.5kb of TSS) of old and young genes of *Arabidopsis thaliana*.

**Supplementary Figure 4:**
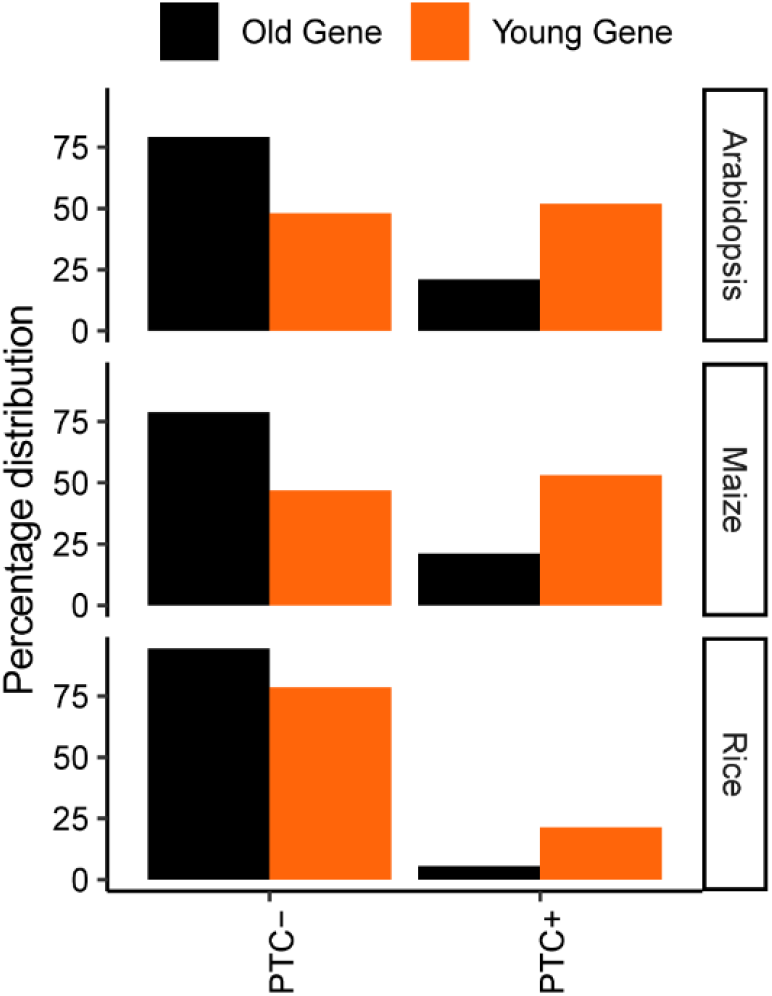
A bar plot showing the distribution of premature translation termination codons (PTC) in old and young genes of Arabidopsis, rice, and maize.

## Notes

### Competing Interest Statement

The authors have declared no competing interest.

